# A Dual-Readout Photonic Sensor for Simultaneous Measurement of Enzyme Activity and Concentration

**DOI:** 10.1101/2025.05.12.653528

**Authors:** Jordan N. Butt, Daniel J. Steiner, Michael R. Bryan, Katie E. Mann, Benjamin L. Miller

## Abstract

Enzyme assays are a cornerstone of basic biology and clinical diagnosis. Typically, enzyme activity is measured, but concentration of the enzyme is also of interest, as are comparisons between concentration and activity. In these situations, separate concentration (i.e. ELISA) and activity (i.e. absorbance) assays are required to fully quantify. Here, we report a multiplex disposable photonic biosensor for simultaneous measurement of enzyme activity and concentration. Capture of the enzyme by a ring resonator-bound antibody produces a red shift in resonance, which can be referenced to a nonspecific binding control. At the same time, enzyme-mediated degradation of a ring-bound substrate produces a resonance blue shift, which can be referenced to a peptide inert to enzymatic cleavage. We tested the dual assay with human Cathepsin-L, dysfunction of which is a hallmark of several diseases, including COVID-19, kidney failure, and cancer. Both assays were found to be well-behaved analytically, with lower limits of detection of 2.0 ng•mL-1 (concentration) and 1.8 ng•mL-1 (activity), well within the range clinically relevant concentrations. Further assessment with a panel of 25 single-donor human serum samples confirmed utility of the assay in a complex, biologically relevant matrix. This approach therefore serves as a useful method for Cathepsin-L detection, and a prototype for other dual-mode photonic enzyme assays.

## Introduction

Quantification of enzymatic activity and enzyme concentration are crucial in research^1^, medicine^2^, biotechnology^3^, and many other areas.^4–6^ While often only one of these measurements is required (activity or concentration), there are other instances where knowledge of both activity *and* concentration is important. For example, in a variety of cancers, enzyme activity is known to change as concentration remains the same.^7^ Likewise, enzyme inhibition is a hallmark of many therapeutics^8^; understanding the efficacy of inhibition is only possible when the concentration of the enzyme being inhibited is found to be constant. Unfortunately, two separate assay formats are currently required for full quantification of an enzyme: a capture-based assay such as ELISA for enzyme concentration, and a separate assay to quantify the enzyme’s activity. This creates obvious disadvantages in workflow complexity, and increases the potential for error. The clinical need for a multimode biosensor able to address both detection mechanisms has been discussed.^9^

While combined assays for enzymatic activity and concentration have not been reported, significant effort has been devoted to the development of novel approaches for each separately. In particular, optical approaches including those based on photonic devices have been widely studied as such devices are readily manufactured^10^, relatively inexpensive^11^, and useful in a wide range of applications^12^. Photonic sensing of enzymes via antibody capture is well known^13^, but photonic structures have also been used for detection of enzymatic activity. For example, color changes in porous silicon have been used to report enzymatic activity^14^. Real-time monitoring of a mesoporous silicon double layer^15^ has been used to sense protease activity, as has enzymatic degradation of gelatin through an optically responsive alumina film^16^. Liquid crystal biosensors have been used to measure enzyme activity in a number of medical and environmental applications^17^. The use of a photonic sensor for real-time measurement of drug-dependent enzyme activity inhibition has been demonstrated^18^. An assay employing single-walled carbon nanotubes has been described for monitoring cholinesterase activity and inhibition in plasma, but the approach is limited to fluorescent substrates^19^.

In order to develop an assay capable of simultaneously quantifying enzyme concentration and activity in a sample, we built on a “disposable photonics” ring resonator platform recently developed in our laboratory.^11,20^ Widely studied over many years by many groups as photonic biosensors^21–27^, a ring resonator consists of a circular waveguide placed near a linear (“bus”) waveguide such that light couples evanescently from the bus waveguide to the ring. Wavelengths of light satisfying the resonant condition (eq. 1; n_eff_ is the effective refractive index of the medium) circulate in the ring and are 180º out of phase with light in the bus waveguide, producing a drop in output power at the resonant wavelength (λ_res_). Since the resonant wavelength is a function of the effective refractive index, molecules of interest binding to the ring via an immobilized capture probe produce a red-shift (increased effective refractive index) in proportion to the amount of material bound. Subtraction of the red-shift observed in a paired ring resonator functionalized with a control probe allows for correction for nonspecific binding and determination of the true binding signal^28^. We hypothesized that enzymatic cleavage of a sensor-bound peptide substrate would result in a blue-shift (decreased refractive index) due to the loss of mass. Combining this with antibody capture of the enzyme on a neighboring ring would enable simultaneous quantification of enzyme concentration and activity in a single test (Figure 1). Example shifts seen during assays are shown in figures S2 (antibody capture of enzyme) and S3 (enzymatic cleavage by the enzyme).

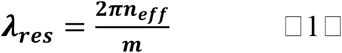

**Figure 1.**
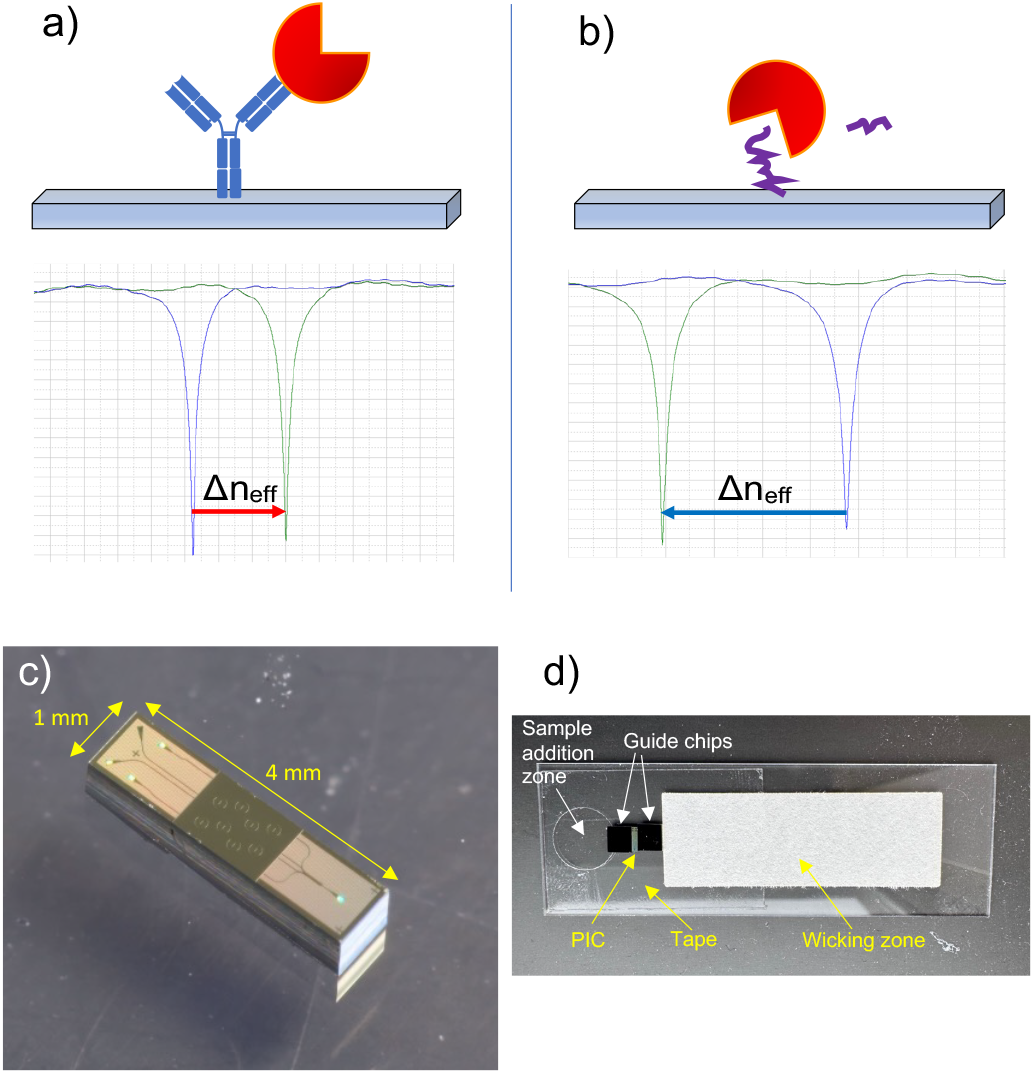
Schematic of sensing concept used in the antibody and enzymatic sensor. a) Capture of enzyme by an immobilized antibody produces a resonance red-shift. b) Cleavage of an immobilized substrate produces a resonance blue-shift. Both assays can operate simultaneously on neighboring rings of a photonic chip. c) 8-ring photonic integrated circuit (PIC) used in these experiments. d) Glass-Laminated Adhesive Microfluidics (GLAM) card used for running assays.

Sample delivery is a critical component of any diagnostic assay. In our previous work, we used injection molded polystyrene micropillar cards to form the microfluidic channel and drive capillary flow of sample to the sensor surface. While effective, this approach relies on precisely molded parts, exact placement of multiple components, and limits sample volumes. Here, we have simplified the microfluidic design to a format we term Glass Laminated Adhesive Microfluidics (GLAM). This approach layers doubled-sided adhesive, in which a channel layer has been defined, onto a glass microscope slide. This provides a simple, repeatable, and effective mechanism for fluid delivery to the sensor chip surface. The sensor chip (Figure 1c) is placed on the channel layer and is flanked by silicon chips to promote uniform flow. Sample volumes for the assay can be tuned by changing the composition and geometry of a reservoir pad that is placed downstream from the sensor chip (Figure 1d).

As an initial test of the dual-format enzyme assay, we focused on Cathepsin L (CTSL). CTSL is a lysosomal cysteine protease, involved in protein processing and in breaking down cells during apoptosis prior to their recycling^29, 30^. Elevation of CTSL in disease relative to baseline levels has been observed in numerous pathologies, including kidney failure^31,32^, parasitic infection^33^, carcinoma^34^, cardiomyopathy^35^, pancreatic cancer^36^, and infection with the SARS-CoV-2 virus, the causative agent of COVID-19^37^. CTSL has also been shown to play a role in the activation of SARS-CoV-2.^38^ Differing levels of CTSL between severe vs. mild/asymptomatic cases of COVID-19 enable a metric for early risk assessment of the infection. The quantifiable CTSL enzymatic activity^39^ has also been shown to be an additional factor in severe cases of COVID-19^37^, as well as being an indicator in numerous stages of cancer^40^. Numerous inhibitors of CTSL enzymatic activity are known;^41^ its inhibition is one of the targets of the commercial drug Paxlovid^42^.

Azocasein, a dye-conjugated derivative of casein, is the standard substrate for photometric CTSL assays,^29^ and was used as the immobilized enzyme substrate for our photonic assay. CTSL activity is typically measured at a pH between 5.5 and 6.5, and specificity for the substrate casein/azocasein is known to be high in this pH range^29,43,44^. We anticipated that ring resonators bearing immobilized casein would show a blue-shift in proportion to the amount of substrate cleaved off the surface of the ring.

## Materials and Methods

Human CTSL and Cathepsin-B expressed from human HEK 293 cells was obtained from Acrobiosystems (Newark, DE). Rabbit anti-human CTSL antibodies were obtained from Sino-biological, Inc (Wayne, PA). Goat anti-fluorescein (anti-FITC) antibody and bovine serum albumin (BSA) used as nonspecific binding and substrate controls were obtained from Rockland Immunochemicals (Limerick, Pennsylvania). E-64 protease inhibitor, casein, and azocasein were obtained from Sigma-Aldrich (St. Louis, MO). Gibco Premium Fetal Bovine Serum (FBS) was purchased from ThermoFisher (Waltham, MA). Assay wash buffer (AWB), which was mixed with FBS at a 4:1 ratio to make FB20 and used to dilute serum samples, consisted of mPBS with 3 mM EDTA and 0.01% Tween-20. The diluent for antibody/antigen printing was modified (i.e., potassium-free) phosphate-buffered saline (mPBS) at a concentration of 0.20 M and a pH of 7.2. (3-glycidyloxypropyl)trimethoxysilane (GPTMS) was obtained from Gelest, Inc. (Morrisville, PA).The diluent for running assays was mPBS, mixed with FB20, at a pH of 5.8, adjusted to ensure CTSL enzymatic activity. Stock PBS was purchased from Sigma-Aldrich and were used as obtained from the manufacturer. The CTSL ELISA kit used to confirm enzyme activity was purchased from RayBiotech (Peachtree Corners, GA). A Perkin Elmer Enspire multimode 96 well plate reader was used for absorbance measurements for both the ELISA and enzyme assays.

Human serum samples were obtained under a protocol approved by the University of Rochester Medical Center Research Subjects Review Board. All subjects were at least 18 years of age at the time of blood draw, and subject to informed consent. Whole blood samples were allowed to clot for 30 minutes after draw. Samples were then spun at 1200 × g for 5 min, and serum was pipetted off into a 15 mL conical tube and spun again for 10 minutes to remove any remaining cellular material. The serum was then aliquoted and stored at −80 °C until use.

### Microring Resonators

The silicon nitride microring resonators used in this work are consistent with our previous designs^11^ but have been adapted to accommodate an increased number of sensing elements within the same chip footprint. The resonators are fabricated with waveguides 1.5 µm wide and 220 nm tall, supporting a single transverse electric (TE) polarization mode. Optical modeling was performed using the finite difference (FD) method in OptoDesigner (Synopsys Photonic Design Suite). A 5.3 µm bottom oxide thickness was used to improve grating coupler performance relative to earlier versions. Each photonic integrated circuit (PIC) includes eight microring resonators, arranged as four channels with two rings per bus waveguide. To accommodate this within the 800 µm-wide usable region of the 1 × 4 mm PIC, the ring diameter was reduced from 198 µm to 164 µm, and the coupling gap was narrowed to 375 nm to compensate for increased bending losses and preserve near-critical coupling. These eight rings are integrated within a 1.3 µm-wide fully etched trench, replacing the previously reported individually etched sensing trenches (Figure S1). This new layout allows for higher integration density and improved fluidic compatibility. Photonic sensors were fabricated using the 300 mm AIM Photonics fabrication line^45^ (Albany, NY). Wafers were then diced by the Aim Photonics Testing and Packaging facility (Rochester, NY).

### Photonic Chip Functionalization

Prior to functionalization, sensor PICs were removed from the wafer and washed for 20 minutes in a 3:1 mixture of sulfuric acid and 25% hydrogen peroxide (“piranha” solution; Caution! Piranha solution is highly caustic and reacts violently with organics). Afterwards, the PICs were washed in nanopure water for 20 minutes before being dried under a stream of nitrogen gas. PICs were next placed in a chemical vapor deposition (CVD) system (Yield Engineering Systems, Fremont, CA) where approximately a monolayer of GPTMS was deposited on the surface with a thickness between 6-10 Å as measured on satellite silicon/SiO_2_ substrates by spectroscopic ellipsometry (J. A. Woolam VASE). Covalent attachment of proteins and peptides to the GPTMS functionalized surface was achieved by spotting directly on the rings using a sciFLEXARRAYER SX piezoelectric microarrayer (Scienion AG, Berlin, Germany), as described previously^46^. The rings used for substrate cleavage were spotted with bovine serine albumin (BSA) as a control and azocasein or casein as a substrate at 1000 µg•mL^-1^. The rings used for antibody capture were spotted with CTSL monoclonal antibodies at 500 µg•mL^-1^ and referenced to anti-fluorescein isothiocyanate (anti-FITC) as a non-specific binding control, spotted at 500 µg•mL^-1^. All rings received approximately 1 nL of the respective solution. The print layout is shown in Supplementary Figure S4. Sensor chips were maintained at 75% humidity overnight to allow reaction with GPTMS to proceed to completion, then used within a day.

### Assay Consumable Assembly

PICs were integrated with a simple, inexpensive glass microscope slide-based microfluidic card (GLAM) designed to provide passive flow of sample liquids into the photonic chip for analysis. 75 × 25 mm glass slides were obtained from VWR (Radnor, Pennsylvania). Glass slides were treated before assembly with UV/ozone for five minutes to increase hydrophilicity. A section of 26 µM thick double-sided adhesive (Adhesives research, York, Pennsylvania) patterned using a laser cutter (Full Spectrum Laser, Hobby Series 20 × 12, Las Vegas, NV) was placed on the treated glass slide. This adhesive layer defines the microfluidic channel geometry and structures a channel height that facilitates efficient analyte delivery. The sensor chip was placed onto the double-sided adhesive such that there was conformal contact of the input and output gratings and the adhesive. The sensor region of the chip is located directly above the open channel. After placing the sensor chip, two silicon chips, also treated with UV/ozone, were placed on the adhesive flanking the sensor chip to stabilize flow. Finally, a strip of filter paper (Q1, Whatman, Little Chalfont, UK) was placed downstream of the PIC, to facilitate continuous flow and provide a waste reservoir for the sample volumes used.

### Apparatus Overview

The optical apparatus used for all photonic measurements follows the methods previously described^11^ and is shown in Supplementary Figure S5. A tunable laser (Keysight 81606A0) is directed through polarization controllers (Thorlabs FPC561 with SMF-28 FC/PC connectors) to maximize transverse electrical (TE) polarization relative to the orientation of the silicon nitride waveguide. Light is directed through the input of the optical hub (15 X 15 X 16 mm; diamond-turned by Syntec Optics, Rochester, NY) and aligned to the grating input. Light from the four output gratings is collected by the output section of the optical hub and directed through a custom fiber bundle (IDIL Optics) of multimode fibers (Thorlabs FP200ERT) to four channels of the optical power meter (Keysight N7745A). Alignment of the PIC is achieved through an IR microscope. A 5× IR objective lens (Mitutoyo Plan Apo NIR 46−402) with on-axis illumination directs light though a long-pass dichroic mirror (Thorlabs DMLP950R) to the IR camera (WiDy InGaAs 650. Proper alignment is confirmed through visual inspection of the IR microscope image, maximization of output power relative to other alignments, and resonance spectra.

The tunable laser and optical power meter are connected to a computer via a general-purpose interface bus (GPIB) and are controlled by the Insertion Loss software of the Keysight Photonic Application Suite (N7700A). Spectral measurements were recorded by repeated wavelength sweeps. Ten-nm spectra were taken continuously at 1 pm resolution, centered on 1550 nm, with each spectral sweep taking approximately 6 s, as previously described^46^. Output spectra were processed automatically through custom software written in Python and deployed on Anaconda, as previously described^11^.

### Sample Preparation and Assay Procedure

To make serial dilution samples, commercially obtained Pooled Normal Human Serum (PNHS) from 2018 was used for determination of detection limits. Pre-COVID-19 pandemic PNHS serum was used in order to avoid potential complications from the inadvertent inclusion of samples from subjects with undiagnosed COVID-19 disease in the pooled serum, which would lead to an unnaturally high baseline. 100 μL of FB20 was mixed with 900 μL mPBS at pH 5.8, or at the same ratio with different volumes, to make the sample diluent, FB2. A 100 µg•mL^-1^ solution of Cathepsin-L was prepared in FB2, then diluted with FB2 in 10-fold increments to produce a dilution series down to a concentration of 100 pg•mL^-1^ in a constant background matrix. These samples were used to generate the serial dilution responses. In all assays, 75 μL of fluid was added to the sample inlet of a GLAM card. Data collection started immediately. Assays were conducted at ambient laboratory temperature (22 ^o^C), and endpoints were measured at 5 minutes. Each sample was run 3 times. The data was then fit to a four-parameter logistic (4-PL) equation in Origin 2023 (OriginLab Corporation, Northampton, MA).

To test the effect of an enzyme inhibitor on the assay, unspiked serum samples used as controls were formed by mixing 90 µL of FB2 with 10 µL of serum. Spiked serum samples were made by mixing 80 µL of FB2 with 10 µL of serum and 10 µL of a 100 ng•mL^-1^ cathepsin-L solution made in the same manner as the serial dilution measurements. Each sample was then vortexed to ensure it was fully mixed. The spiked sample was then split into an additional two samples, and 1 mg of the E-64 inhibitor was added to one of the two samples. Each sample was then vortexed to ensure it was fully mixed. Samples were allowed to incubate at 4 ^o^C for 3 hours prior to assay, and then run immediately. To test potential interference, 80 µL of FB2 with 10 µL of serum and 10 µL of a 100 ng•mL^-1^ cathepsin-B solution, made in the same manner as the cathepsin-L solutions.

To assess the effect of background matrix variability, 25 single-donor serum samples were selected from individuals between age 18 and 35 without any known health conditions. Each sample were split into spiked and unspiked samples. Unspiked serum samples were formed by mixing 90 µL of FB2 with 10 µL of serum. Spiked serum samples were made by mixing 80 µL of FB2 with 10 µL of serum and 10 µL of a 100 ng•mL^-1^ cathepsin-L solution made from the serial dilution measurements. Each sample was then vortexed to ensure it was fully mixed. As above, each sample was allowed to incubate at 4 ^o^C for 3 hours prior to assay.

## Results

### Serial Dilution Curve Generation for Cathepsin-L

The concentration and enzymatic activity of commercial CTSL used throughout this study were confirmed using reference-standard ELISA and UV-Vis enzymatic assays (Supplementary Information). The reference UV-Vis enzymatic assay was likewise used to establish the baseline concentration and activity of PNHS and FBS prior to doping in recombinant CTSL, and to confirm that cathepsin-B was inactive in this matrix (Supplementary Information Figures S6, S7). The presence of Cathepsin-L in PNHS was confirmed with an ELISA assay, and found to be 1.97 ng•mL^−1^, and the spiked serum had a net concentration of 11.97 ng•mL^-1^. This encompasses the range of reported healthy and elevated levels of Cathepsin-L in serum^36^. Dilution series of CTSL in PNHS accounted for the baseline 1.97 ng•mL^-1^ concentration.

For photonic assay serial dilution samples, the limit of blank (LOB) was determined and added to the standard deviation of low-concentration samples to derive lower limits of detection (LLOD) of human CTSL antibody capture and enzymatic activity.^47^ Results are displayed in Figures 2 and 3. Both assays are well-behaved, with calculated LLOD of 2.0 (substrate cleavage) and 1.8 ng•mL^-1^, respectively. Both are below the required LLOD for CTSL in clinical situations of 2.2 ng•mL^-1 37^. Because this assay is label-free, its dynamic range may be tuned based on modifications to the enzyme substrate: we observed that immobilized casein produced a lower magnitude shift in the enzymatic assay (due to both its lower mass and reduced refractive index change on cleavage) relative to the dye-tagged azo-casein (Supplementary Figure S8).

**Figure 2.**
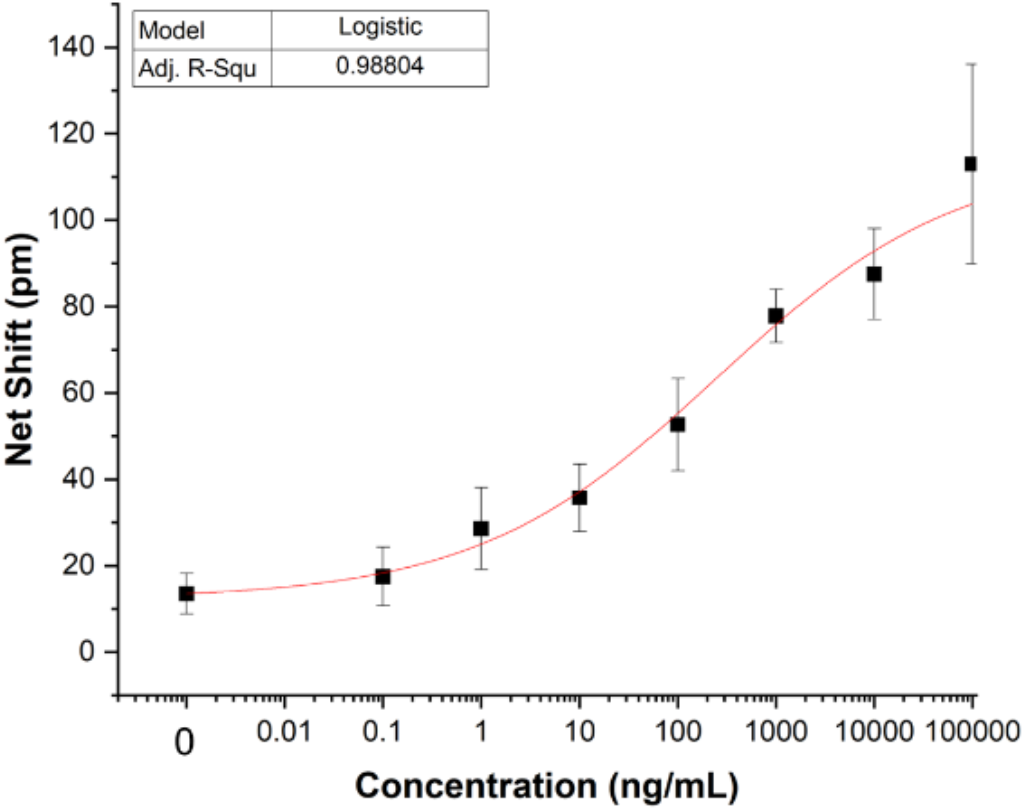
Serial dilution of Cathepsin-L (antibody capture assay). Each point represents the average of 3 replicate assays, with error bars ± one standard deviation shown. Red line shows nonlinear least squares fit of the data to a logistic model. Calculated LLOD = 2.0•ng•mL^-1^.

**Figure 3.**
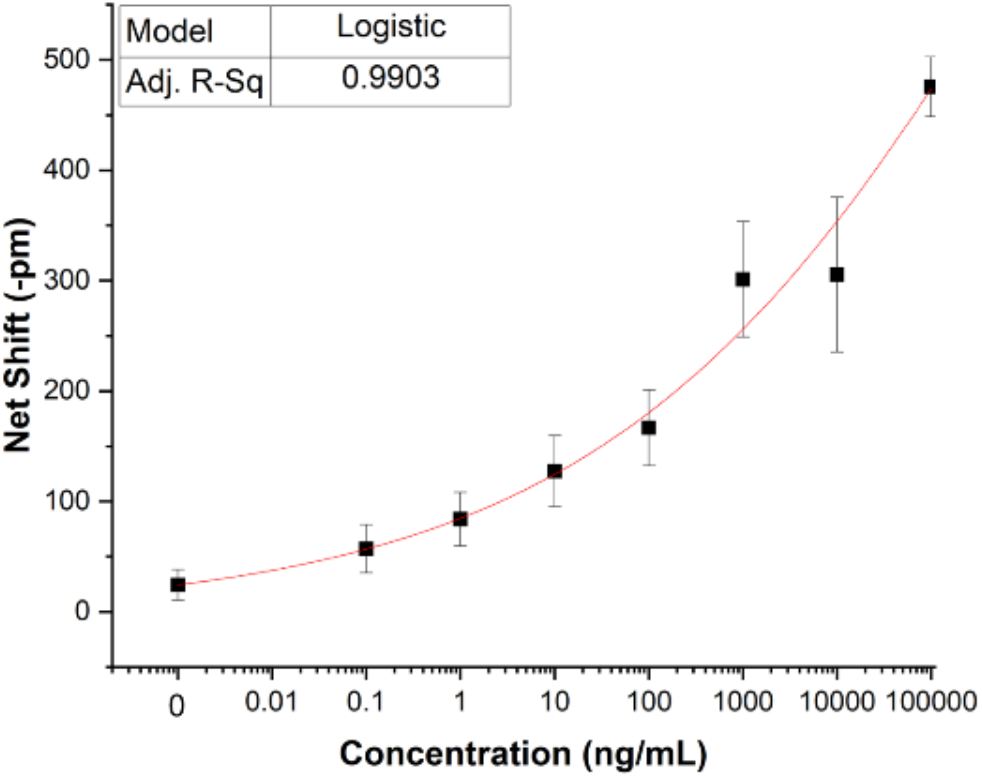
Serial dilution Cathepsin-L response (enzymatic degradation of immobilized azocasein). To enable curve fitting, the absolute value of the net shift is used, instead of the negative shift. Each point represents the average of 3 replicate assays, with error bars ± one standard deviation. The red line shows the calculated logistic fit of the data. Calculated LLOD = 1.8 ng•mL^-1^.

### Dual Antibody and Enzymatic Assay using Pooled Human Serum

To assess the ability of the dual assay to report changes in CTSL concentration and activity, we compared responses of 1:10 diluted PNHS, PNHS + 10 ng•mL^-1^ CTSL, PNHS + 10 ng•mL^-1^ cathepsin-B, and PNHS + 10 ng•mL^-1^ CTSL + E-64, an irreversible inhibitor of cysteine proteases^48^. The baseline measurement of CTSL concentration and CTSL activity present in PNHS and PNHS + 10 ng•mL^-1^ CTSL (Figure 4) were consistent with measurements done in the context of the serial dilution experiments. The addition of the inhibitor agent changes the enzyme activity to a point where the only observed shifts are within the noise, while the unchanged concentration results in antibody capture that is similar to the sample without an inhibitor agent. These results highlight a key advantage of this sensor, as the change in concentration and enzyme activity can be measured in a single test. T-tests were performed to ensure statistical significance. All differences were significant at 95% confidence, except the difference between the antibody capture for the spiked and spiked + inhibitor solutions. Supplementary Table T1 details the statistical results.

**Figure 4.**
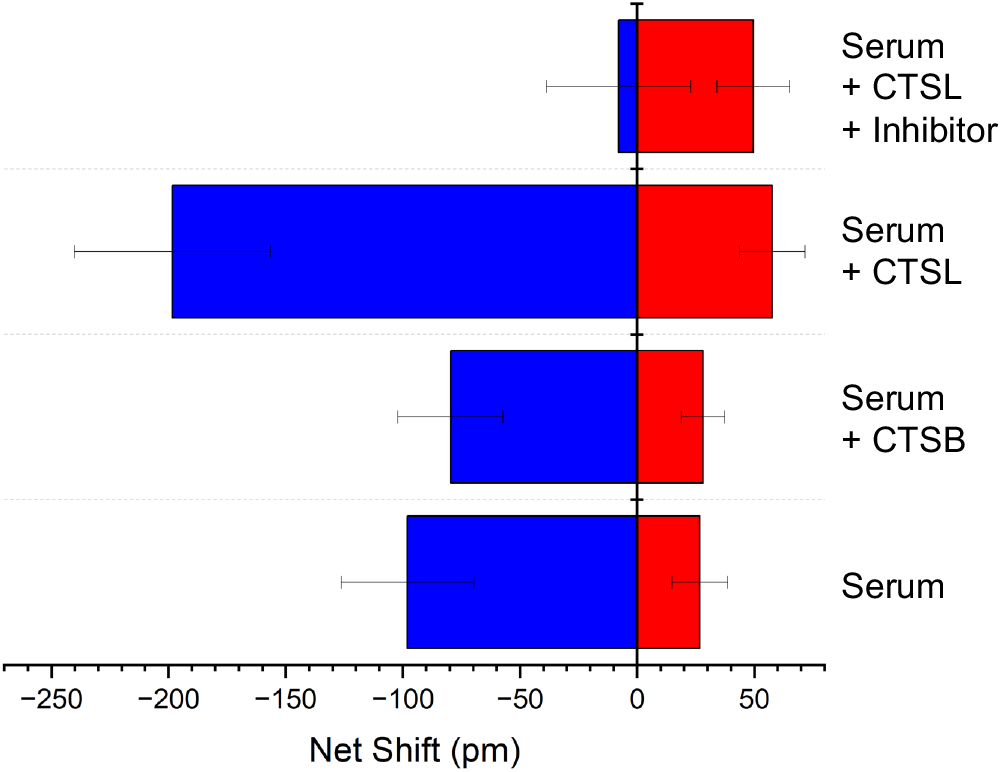
Dual photonic assay successfully reports changes due to changes in CTSL concentration and inhibition of CTSL activity in human serum. Baseline concentration and activity of pooled normal human serum (PNHS) alone, doped with 10 ng•mL^-1^ CTSL, and doped with 10 ng•mL^-1^ CTSL plus inhibitor were recorded. Serum doped with 10 ng/mL CTSB produces a response statistically indistinguishable from that produced by serum alone, demonstrating the specificity of the assay. Each value represents the average of 6 replicate measurements, with error bars plus/minus one standard deviation.

While the specificity of Cathepsin-L for azocasein at acidic pH is well known^44^, we confirmed this was the case in our assay as well by comparing the response to samples doped with Cathepsin-B. No statistically significant difference in response was observed for PNHS doped with 10 ng/mL CTSB, vs. just the serum alone (Figure 4).

### Dual Antibody and Enzymatic Assay Single Serum Samples

Finally, we evaluated the performance of the dual CTSL assay in 25 single-donor human serum samples. Baseline responses were first measured for each sample; 10 ng•mL^-1^ CTSL was then doped into the sample, and it was re-measured. Figure 5 shows the differential response for all 25 single-donor serum samples after addition of 10 ng•mL^-1^ CTSL relative to baseline. We were pleased to observe that the assay provided relatively consistent responses, with a coefficient of variation (CV) of 16.4 % for the enzymatic cleavage and 15% for antibody capture. When not incubated at consistent times, we observed that variation increased significantly, as would be expected for any enzyme assay.

**Figure 5.**
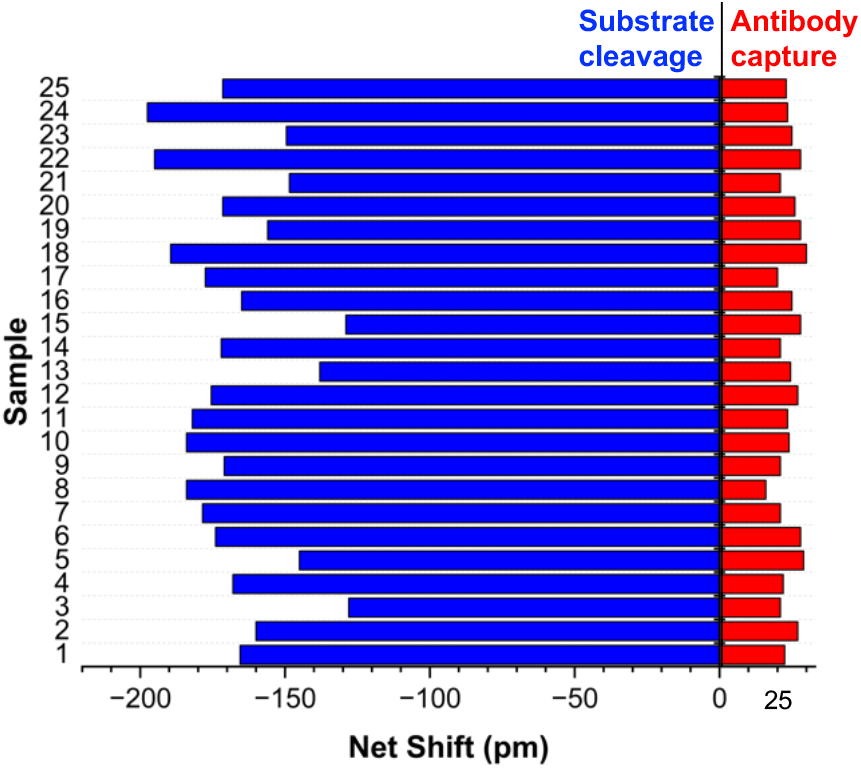
Differential response for 25 single-donor human serum samples assayed before and after being doped with 10 ng•mL^-1^ CTSL.

## Discussion

A central advantage of label-free sensors is the ability to build in multiplex sensing capability without increasing complexity of the assay. Beyond this, an important but relatively unstudied area is that of multiplexing different types of assays, i.e. multimodal sensing, to increase the different types of information that can be obtained from a system under study. Here, we demonstrate a multiplex photonic sensor capable of simultaneously quantifying enzyme activity and concentration in a sample. Both detection modes are analytically well behaved, specific, and usable in the complex and variable background matrix of human serum. Importantly, the assay was able to distinguish inhibition of CTSL by E-64 inhibitor from a decrease in concentration of the enzyme.

We observed that tuning both the mass and refractive index of the immobilized substrate may be used to alter the performance of the assay in subtle ways, by comparing responses with casein and azocasein. Casein has a refractive index of 1.5749, whereas the dyed azocasein has a refractive index of 1.850. Thus, cleavage of azocasein results in a larger change in effective refractive index than casein. This provides intriguing possibilities for assay development.

In the clinical context, as already discussed, CTSL is a useful marker for severe COVID-19^37^, and an essential enzyme in the SARS-CoV-2 infection pathway^38^. A single study noted 42 different clinical enzymes that diagnose various disorders, including cancer, liver failures, heart attacks, and more^51^. Many of these should be amenable to the assay format we report here. Moving beyond potential clinical applications, the development of enzyme inhibitors is of significant importance,^52^ and the assay format we report here should have utility in that regard. Other potential applications include rapid screening of substrate specificity, both in biomedically relevant contexts and in biomanufacturing.

We note that this approach is not suitable for assaying activity of all enzymes: requirements include that the enzyme have a substrate that (1) may be immobilized on the surface of the sensor without compromising the ability of the enzyme to react with it, and (2) the enzymatic reaction must cause a significant change in mass (either increase or decrease) of the sensor-immobilized substrate. Examples of enzymes and substrates potentially fitting these criteria include collagenase (enzymatic cleavage of collagen; loss of mass) and ubiquitin ligases^53^ and artificial peptide ligases^54^ (enzymatic conjugation of peptides; increase in mass). Efforts to test these and other systems are underway in our laboratory.

## AUTHOR INFORMATION

### Corresponding Author

Benjamin L Miller;

### Author Contributions

The manuscript was written through contributions of all authors. All authors have given approval to the final version of the manuscript. **Jordan Butt:** Conceptualization, Methodology, Investigation, Validation, Formal Analysis, Writing-Original draft preparation, Visualization. **Daniel Steiner:** Conceptualization, Methodology, Validation. **Michael Bryan:** Software, Investigation, Methodology. **Katie Mann:** Investigation, Validation **Benjamin Miller:** Conceptualization, Methodology, Writing-Original draft preparation, Writing-Review and Editing, Supervision, Funding acquisition.

## ACKNOWLEDGMENT

The authors thank Harold Warren and Raymond Jakubowicz for helpful discussions during the preparation of this manuscript.

## ABBREVIATIONS

CTSL: Human Cathepsin-L
GLAM: Glass Laminated Adhesive Microfluidics
BSA: Bovine Serum Albumin
PNHS: Pooled Normal Human Serum

## TOC Graphic

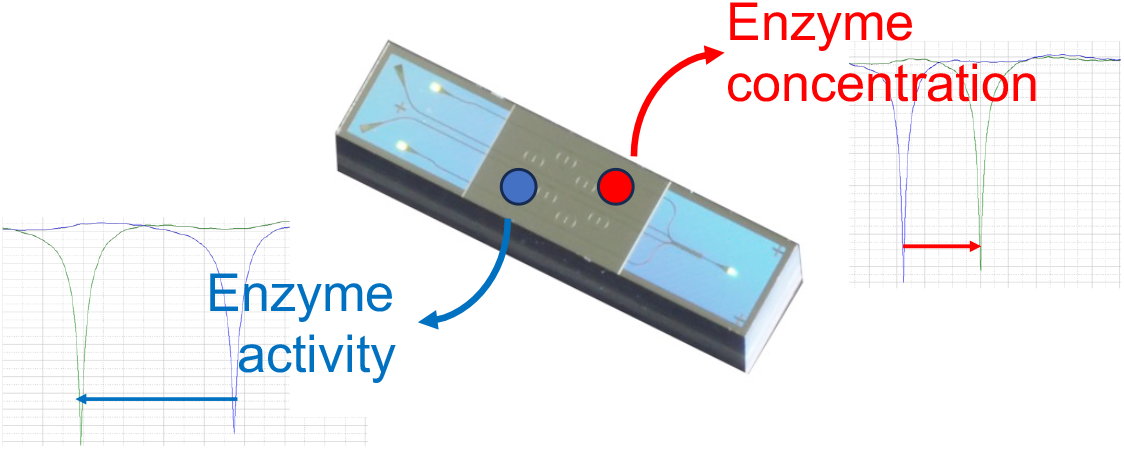

