## Supplementary Information for "A Dual-Readout Photonic Sensor for Simultaneous Measurement of Enzyme Activity and Concentration"

Table of Contents

Comparison of individual trench and full trench photonic sensors…………………..…S2

Example of antibody capture response…………………………………….……………S3

Example of substrate cleavage response………………………………….……………S3

Sensor print and flow orientation schematic…………………………….……………..S4

Schematic of optical apparatus……………………………………………….………… S5

ELISA assay ………………………………………………………………………….……...S6

UV/Vis absorbance enzymatic assays……………….…………………………...……S7

Comparison of enzymatic substrates...............................……………………….…..S8

Statistical tests for sera ± CTSL, CTSB, E-64 inhibitor...………………….……………S9

References………………………………………………………………………………S9

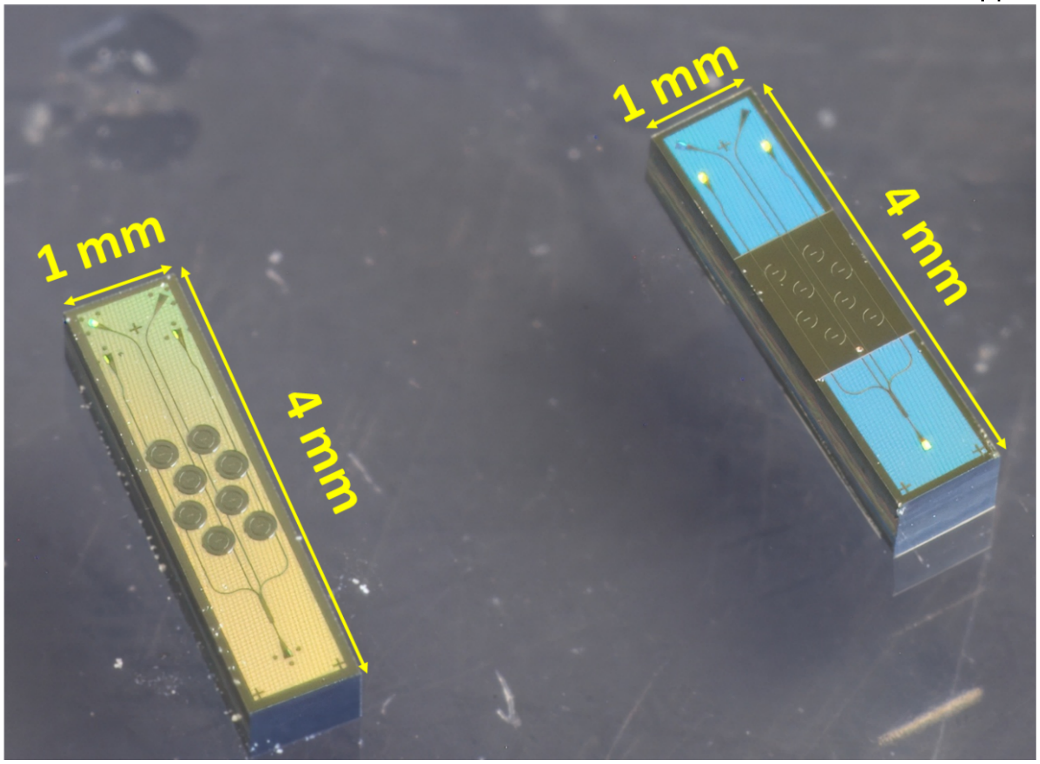


**Figure S1**. Comparison of a PIC with individual trenches around each ring resonator (left) and a PIC with a unified trench (right; used in this work).

**Time-dependent sensor response**

A representative example of the reference-subtracted signal seen for anti-CTSL antibody capture is shown in Figure S2. For enzymatic assays, CTSL-mediated substrate cleavage results in a blue shift (Figure S3).

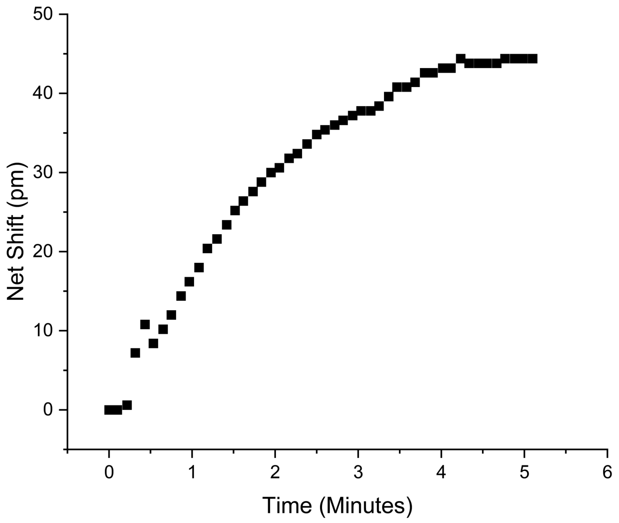


**Figure S2**. Representative red-shift response as a function of time seen from antibody-mediated capture of CTSL on a ring resonator. Concentration: 1000 ng/mL in pH 5.8 mPBS with FB20.

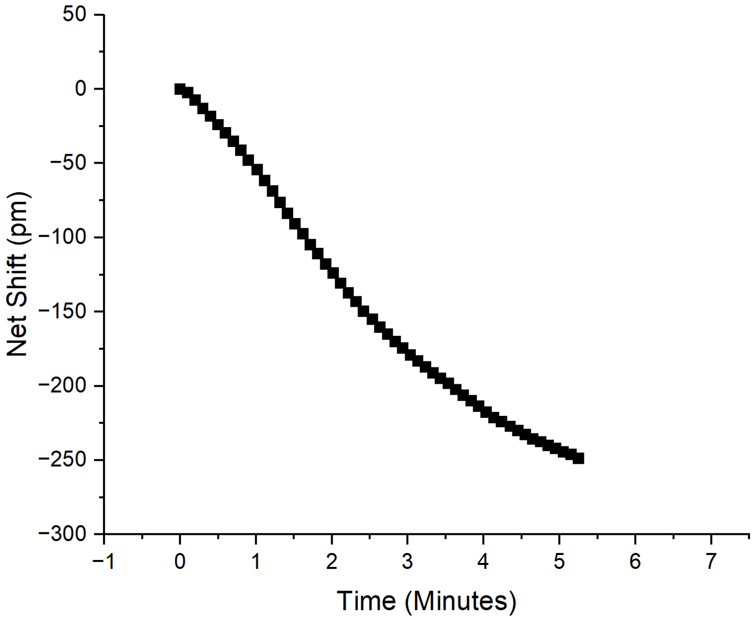


**Figure S3.** Representative time-dependent blue shift seen from substrate cleavage by CTSL of azo-casein on a ring resonator. Concentration: 100 ng/mL in pH 5.8 mPBS with FB20.

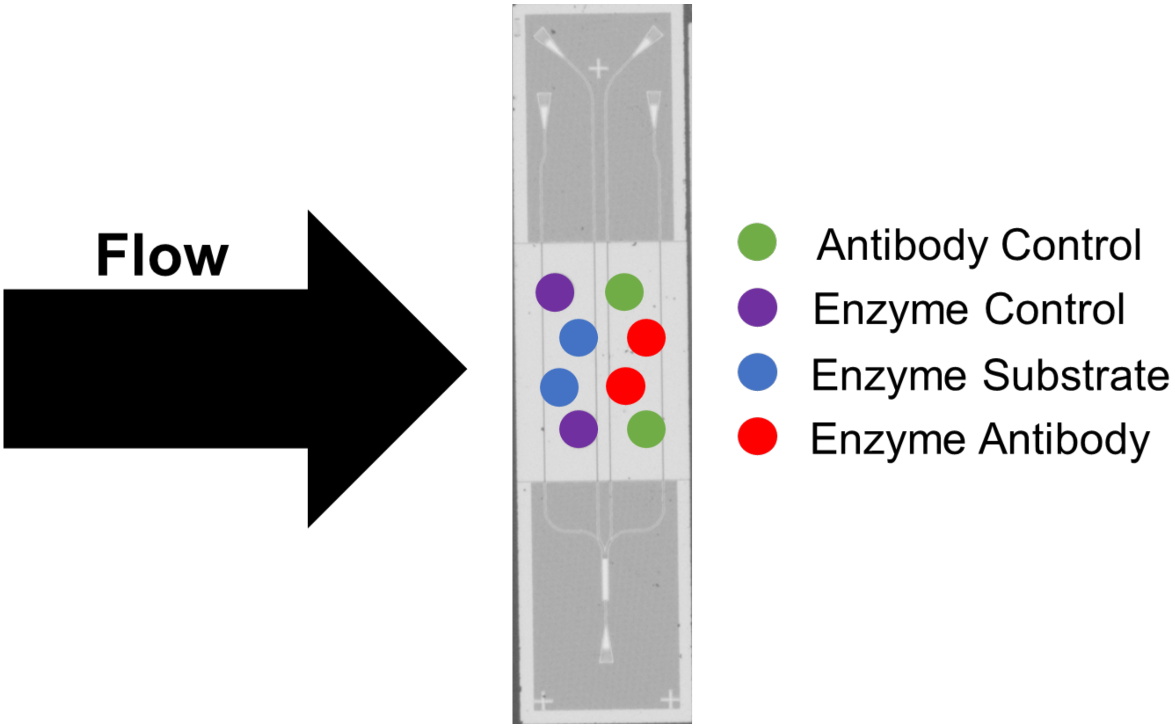


**Figure S4**. Schematic of probe and substrate print geometry, along with orientation of the PIC relative to the direction of microfluidic flow after assembly into a Glass Laminated Adhesive Microfluidics (GLAM) card.

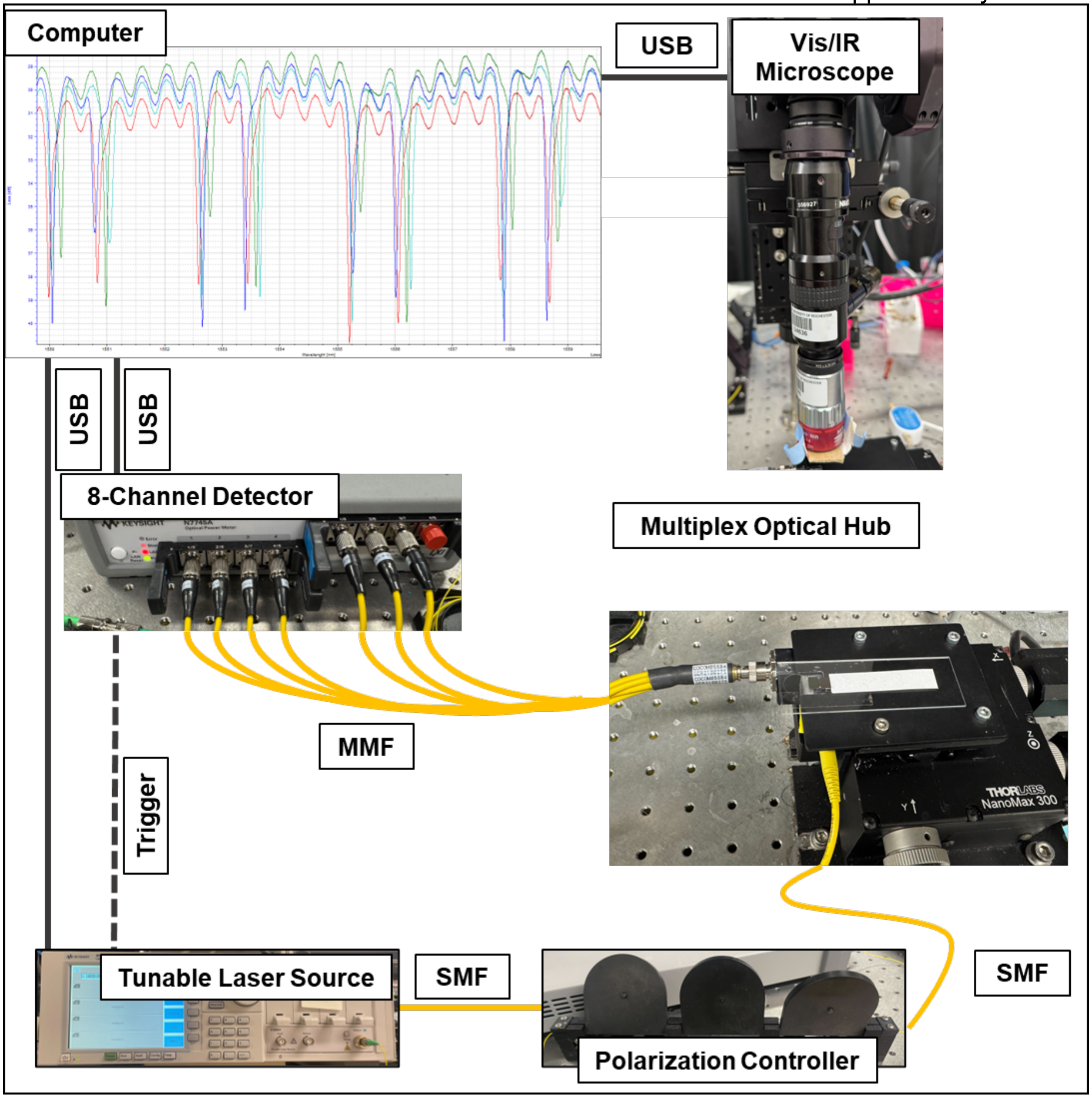


**Figure S5**. Schematic representation of the multiplex optical hub photonic biosensing apparatus. Vis, visible-wavelength light; IR, infrared-wavelength light; SMF, single-mode fiber; MMF, muti-mode fiber; USB, universal serial bus. A GLAM (Glass Laminated Adhesive Microfluidics) disposable card is shown resting on the optical hub stage.

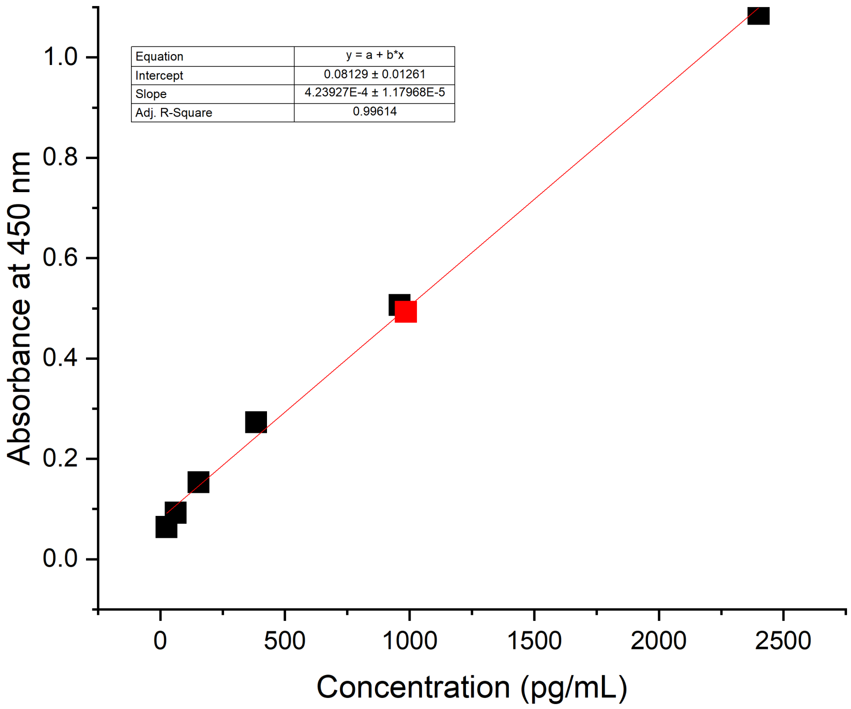


**Figure S6**. ELISA assay for cathepsin-L. Black squares are known concentrations of cathepsin-L in buffer; red square is from the commercial pooled normal human serum, with the measured absorbance used to extract the concentration with the known ELISA dilution response. The ELISA assay had a linear range specified from 24 pg•mL$^{-1}$ to 6 ng•mL$^{-1}$, within the expected range of CTSL in serum.

**Confirmation of Cathepsin-L concentration and activity**

To confirm enzymatic activity of Cathepsin-L, 200 µL of 0.2 mg $mL^{-1}$ azocasein in mPBS pH = 5.8 was mixed with 100 µL of cathepsin-L at varying concentrations. A UV-vis absorption peak of the azocasein was taken and a peak was identified at 440 nm. This is consistent with the information provided by the supplier[1], although the absorbance peak of azocasein can change due to the conjugated dye[2]. The absorbance was then measured at 440 nm every five seconds for one minute. The resulting shifts in absorbance were then analyzed, with the endpoint shift in absorbance to concentration. The $V_{max}$ can be extracted from these data by the molar absorbance of azocasein, which was not calculated because the supplier provided a range of values for the extinction coefficient[1]. This UV-vis absorbance assay confirms that the concentrations of cathepsin-L used in these experiments exhibit enzymatic activity within a linear range.

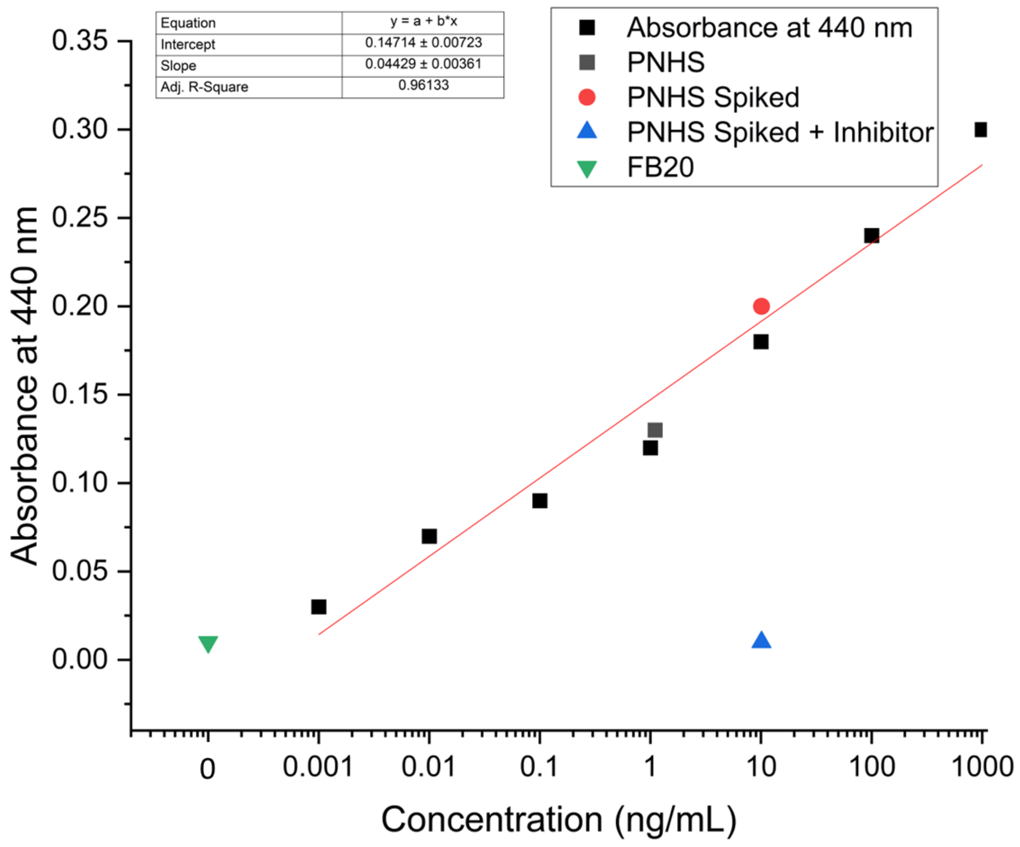


**Figure S7**. Reference standard UV-Vis enzymatic assay used to establish activity of CTSL used in this work, and assess levels of CTSL in PNHS and FB20.

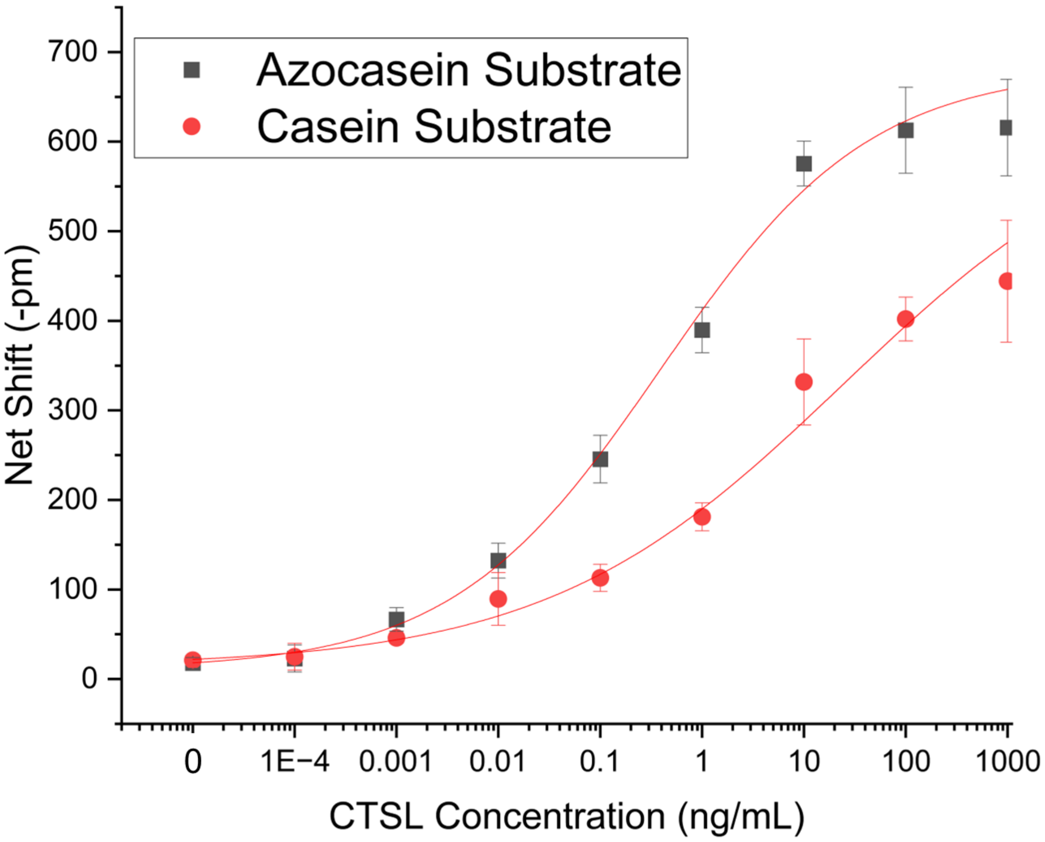


**Figure S8**. Comparison of resonance shifts observed between CTSL substrates azocasein and casein. This assay was performed using a previous-generation micropillar microfluidics card described in Bryan, Butt, et al[3]. Azocasein shows better dynamic range at the lower concentration, while casein shows better dynamic range at higher concentration.

| Solutions compared | A or E? | t-statistic | Probability |
|---|---|---|---|
| U vs S | A | 3.16 | 0.0160 |
| U vs S | E | 2.32 | 0.0487 |
| U vs B | A | 0.185 | 0.857 |
| U vs B | E | 0.959 | 0.369 |
| U vs I | A | 0.719 | 0.495 |
| U vs I | E | 3.19 | 0.0242 |
| S vs I | A | 4.91 | 0.00117 |
| S vs I | E | 7.37 | 0.000154 |
| S vs B | A | 3.35 | 0.0122 |
| S vs B | E | 4.51 | 0.00404 |
| I vs B | A | 2.36 | 0.0457 |
| I vs B | E | 4.78 | 0.00100 |

Table T1. Statistical tests comparing dual assays run on 3 different serums. A = antibody capture assay, E = enzymatic cleavage assay. U = unspiked (PNHS as-obtained), B = PNHS spiked with CTSB, S = PNHS spiked with CTSL, and I = PNHS spiked with CTSL and inhibitor solution.
